# Spontaneous ectopic head formation enables reversal of the body axis polarity in microscopic flatworms

**DOI:** 10.1101/2025.05.14.653983

**Authors:** Katarzyna Tratkiewicz, Ludwik Gąsiorowski

## Abstract

In most of the animals the antero-posterior axis is specified during early embryogenesis. However, in the organisms that undergo somatic asexual reproduction constant re-establishment of the body axis occurs during each asexual act in the context of the fully formed adult body. In microscopic flatworms from the genus *Stenostomum* the new head and tail structures are inserted in the pre-existing body plan during the asexual process known as paratomy. Here, we report a spontaneously occurring developmental error that results in the formation of the worms with double heads at the opposite ends of their bodies that misses the posterior pole identity. In the set of experiments, we show that the double head phenotype is not heritable on the organismal level. Worms originating from the sectioning or fission of the double head animals give rise to the healthy populations that do not display the erroneous asexual development. We also demonstrate that the posterior piece of the double head worm can survive, regenerate the tail on its previously anterior pole and resume the asexual reproduction. Effectively, such regeneration allows reversal of the body axis polarity without impairment of the survival or reproductive abilities of the animal, an exceptionally rare phenomenon among bilaterians.

## Background

In most of the animals the specification of the body axis occurs early on during embryogenesis and represents one of the crucial developmental events, during which polarity of the adult animal is established [1–3]. However, in the organisms that engage in the somatic asexual reproduction, body axes of new individuals have to be specified within the context of the fully formed adult body, either by establishment of the new axes or by developing new individuals along the already established polarity [4–14]. There are three basic types of somatic asexual reproduction, that differ in regards to the alignment of the body axes of maternal organisms and their asexual progeny as well as the sequence of the formation of new tissues during the asexual process. The most widespread type of asexual reproduction is budding, in which new individuals (zooids) bud off from the maternal organism with their body axes specified *de novo* and not aligned with the axis of the maternal animal [4–9, 15–17]. In contrast to budding, the body axis of the maternal individual and the newly formed zooids are aligned and continuous in two other modes of somatic asexual reproduction, architomy and paratomy. The main difference between those types is that in case of architomy the maternal organism first divides, and then regenerates missing body parts, while in paratomy first the new structures of the zooid are formed within the maternal organism, and only later the progeny separates [17–19].

In flatworms (Platyhelminthes), the asexual reproduction evolved many times independently, resulting in different developmental modes used by particular groups [17, 20]. Both paratomy and architomy are attested among platyhelminths, the former in groups Catenulida and Macrostomorpha, and the latter in planarians [17, 18, 20]. Any somatic growth of flatworms depends on the pluripotent stem cells [21–26], that also take part in the process of asexual reproduction [14, 22, 27, 28]. Under normal circumstances differentiation of the flatworm stem cells is controlled by the antero-posterior (AP) gradients of signaling molecules that precisely control axial identity of the developing structures [29–32], even in those groups that do not engage in the asexual reproduction [33, 34]. Concordantly, the experimental disturbance of those signaling molecules can lead to the formation of ectopic head or tail tissues [31, 32, 35, 36]. In planarians, that can reproduce asexually through architomy, the development of surplus heads during regeneration process can also occur spontaneously [37–39] or as a result of extensive dissections [40, 41]. Here, we report that spontaneous ectopic head formation also takes place in a microscopic flatworm, *Stenostomum brevipharyngium*, an emerging catenulid model species, that reproduces through paratomy [22, 42, 43]. In series of experiments, we set out to test viability of the individuals with the ectopic heads as well as their potential for regeneration and asexual reproduction. These results shed light on the extreme developmental plasticity of *Stenostomum*.

## Methods

### Animal husbandry

*Stenostomum brevipharyngium* cultures were ordered in 2010 from Connecticut Valley Biological Supply as *Stenostomum* sp. and since then maintained in the laboratory. The species identification followed the key to the North American species of *Stenostomum* [44]. The cultures were maintained in standardized Chalkley’s Medium (CM) at 20°C in the dark and fed *ad libitum* with the unicellular eukaryote *Chilomonas paramecium*. Under those conditions the animals reproduce exclusively asexually and do not form gonads.

### Animal fixation

For both antibody staining and RNA in situ hybridization we used the same fixation procedure. The animals were first anesthetized in 1.44% (w:v) MgCl_2_ in CM for ca. 10 min. When the worms stopped active movements, they were fixed in 4% (v:v) formaldehyde in PBS + 0.1% Tween-20 detergent (PTw) for 30 minutes. Then the animals were washed three times in PTw and either stored at 4°C (for antibody staining) or washed in ultrapure deionized water, dehydrated in 100% methanol and stored in fresh 100% methanol in −20°C (for RNA in situ hybridization).

### Antibody staining

The animals were washed three times in PBS + 0.1% bovine serum albumine + 0.1% Triton X (PBT) for 15 minutes at room temperature (RT) and then incubated for 30 minutes in 5% normal goat serum dissolved in PBT (PBT+NGS), also at RT. Next, we incubated worms overnight (ON), at 4°C in primary antibodies dissolved in PBT+NGS. We used the following primary antibodies: mouse anti-tyrosinated tubulin, Sigma T9028 (dissolved at 1:500); and rabbit anti-serotonin (5HT), Sigma S5545 (dissolved at 1:250). On the next day, the worms were washed six times with PBT for 15 min and incubated for 30 min in PBT+NGS, both at RT. The animals were subsequently incubated ON at 4°C in secondary antibodies dissolved in PBT+NGS. We used the following secondary antibodies: goat anti-mouse, conjugated with Alexafluor488, Thermo Fisher A-11001; and goat anti-rabbit, conjugated with Alexafluor647, Thermo Fisher A-21244 (both at the concentration 1:250). After incubation in secondary antibodies all steps were performed in RT. The animals were washed three times in PBT, two times in PTw, and incubated for 40 min in Hoechst 33342 dissolved in PTw (1:5000). Then we rinsed the animals twice in PBT, and incubated for 1h in Phalloidin, conjugated with Alexafluor555, Thermo Fisher A34055 (10U/ml dissolved in PBT). Finally, the animals were washed in PBS, mounted in FluoromountG (Thermo Fischer, 00-4958-02), and left overnight at 4°C for tissue clearance and hardening of the mounting medium. The mounted specimens were investigated with the Olympus IX83 microscope with a spinning disc Yokogawa CSUW1-T2S scan head (asexual worms in paratomy) or with the Nikon A1R MP confocal laser scanning microscope (double head worms).

### RNA in situ hybridization

For fluorescent RNA in situ hybridization chain reaction (HCR) v3.0 [45] we used the same DNA probe oligo pools as in [22] ordered at the Integrated DNA Technologies. For in situ staining we used the same protocol as in [46], the worms were rehydrated in Me-OH/PTw series at RT, washed four times in PTw, and then prehybridized in hybridization buffer for 40 min at 37°C. Next, the animals were placed in HCR probe mixtures at 1uM concentration in hybridization buffer and incubated ON at 37°C. On the next day, the probes were removed with four washes of probe wash solution (each 10 min at 37°C), followed by three washes in 5xSSC+0.1% Tween-20 at RT. Then, the worms were incubated in amplification buffer for 30 min at RT. In the meantime, the HCR hairpins (Molecular Instruments: B1H1-546, B1H2-546, B3H1-647, B3H2-647) were prepared by heating for 1min 30sec at 95°C and then cooling at RT in the dark for 30 min. The hairpins were mixed in amplification buffer at 40nM concentration and added to the samples. The animals were incubated in the amplification buffer with hairpins ON at RT in the darkness. On the next day, the samples were rinsed three times with 5xSSC+0.1% Tween-20, twice with PTw and incubated for 40 min in Hoechst 33342 in PTw (1:5000). The stained specimens were washed once in PBS, mounted in Fluoromount G (Thermo Fischer, 00-4958-02), left ON at 4°C and then imaged with the Nikon A1R MP confocal laser scanning microscope.

### Regeneration experiments

For the regeneration experiment the worms were anesthetized with 1% (w:v) MgCl_2_ in CM for ca. 10 min. The worms were cut under the dissecting scope with an eyelash and immediately transferred to a fresh CM. The regenerates were washed few times with CM, to remove the anesthetic medium. Single regenerates were then put into 400μL of CM and kept in darkness at 20°C with addition of 100μL of concentrated culture of *Chilomonas paramecium* (if not stated otherwise). The live regenerating worms were briefly anesthetized with 1% (w:v) MgCl_2_ in CM, imaged on Keyence VHX-7000 Digital Microscope, and then put back into fresh CM.

### Quantification and statistical analyses

For quantification of the regeneration success, we first assign each worm to one of the categories, that linearly described severity of the observed outcome: dead (0), dying (1), retarded regeneration (2), regular non-reproductive (3), regular asexual (4). Next, we used the two-sided Mann–Whitney U test to probe for the difference in distribution of the outcomes between control and experimental animals. To statistically test for differences in asexual reproduction between experimental groups, we calculated the slopes of the reproduction curves for single individuals in each condition (Table S1) and compared them to the slopes of the reproduction curves of the control worms using the two-sided Mann–Whitney U test. For both analysis we defined significance at the level of p-value<0.05.

## Results and discussion

### Characterization of the double head phenotype

*Stenostomum brevipharyngium* is a microscopic flatworm, that can be easily reared in the laboratory [22, 43]. Under favorable conditions *S. brevipharyngium*, similarly to the other species of the genus, engages in asexual reproduction by means of paratomy [14, 22, 43, 47–49]. During the process, the long, well-fed individuals start to produce new anterior structures in the middle of their trunks, in the area referred to as the fission zone. First, the pluripotent stem cells start to divide and accumulate at the lateral sides of the gut, and later they differentiate into new head and pharyngeal tissues [27, 43, 48]. At the late stage of paratomy, the anterior structures of the new individual are fully formed in the middle of the A-P axis, giving rise to the chain, composed of two zooids, which are aligned along A-P axis and attached head-to-tail at the fission zone (Fig1 a–c). In this advanced stage the new, second zooid is equipped with pharyngeal and brain tissues that already resemble those of its maternal individual, both in terms of morphology and polarity (Fig. 1 d–i).

**Fig. 1.**
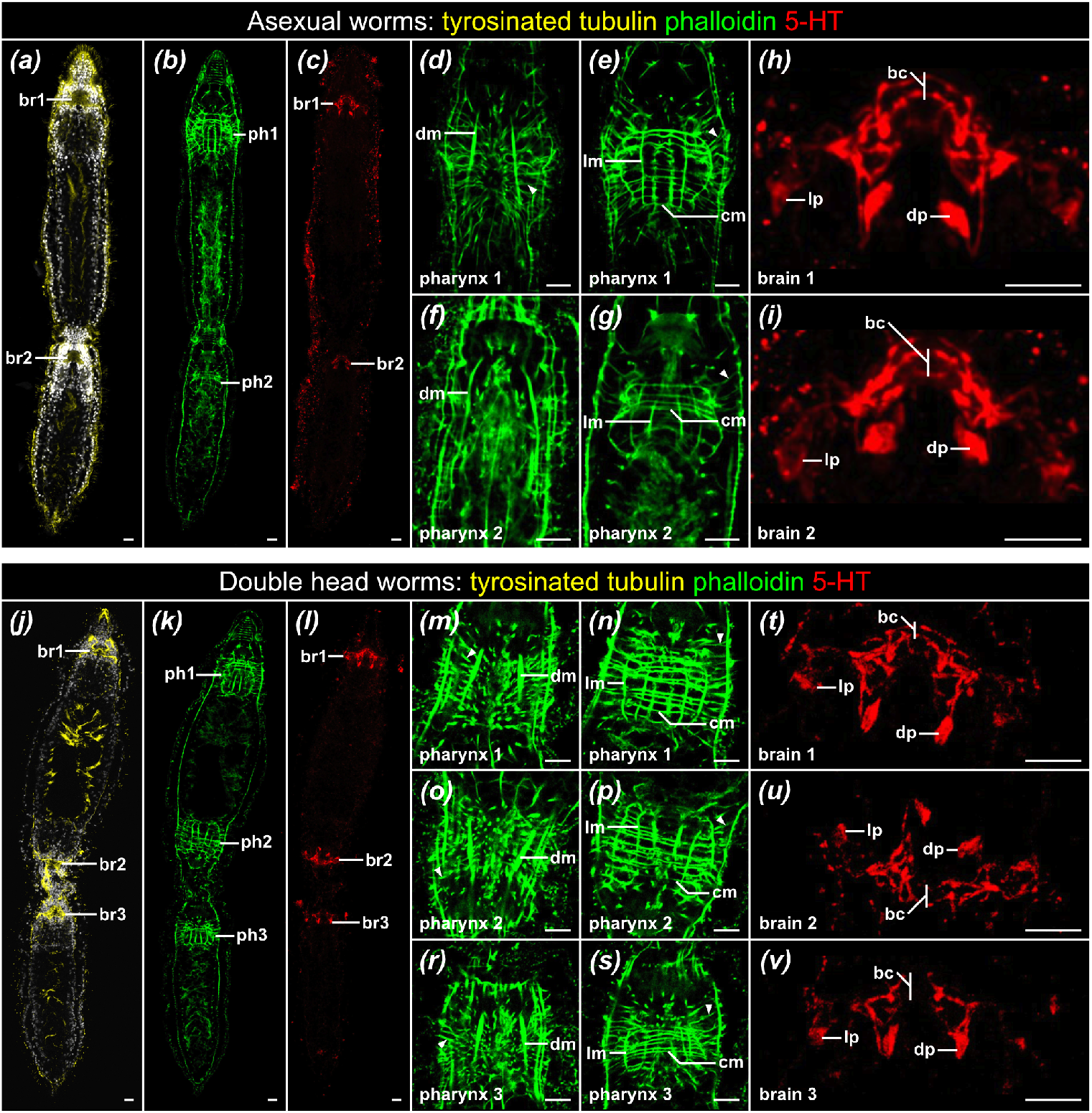
Morphological comparison of the wild type asexual and double head *Stenostomum brevipharyngium*. **(a–c)** General morphology of in the worm at the advanced stage of paratomy. **(d–g)** Detailed morphology of the pharynges in the asexual worm, showing the dorsal pharyngeal region **(d, f)** and the pharyngeal bulb **(e, g). (h, i)** Detailed morphology of the serotoninergic brain in the asexual worm. **(j–l)** General morphology of the double head worm. **(m–s)** Detailed morphology of the pharynges in double head worm, showing the dorsal pharyngeal region **(m, o, r)** and the pharyngeal bulb **(n, p, s). (t–v)** Detailed morphology of the serotoninergic brain in the double head worm. Abbreviations: *bc*, brain serotoninergic commissure; *br1–br3*, brains 1–3; *cm*, circular pharyngeal muscles; *dm*, dorsal pharyngeal muscles; *dp*, dorsal serotoninergic perikaryon *lm*, longitudinal pharyngeal muscles; *lp*, lateral serotoninergic perikaryon; *ph1–ph3*, pharynges 1–3. White arrowheads indicate radial pharyngeal dilator muscles. Scale bars indicate 10 μm.

In one of our routinely run laboratory cultures of *S. brevipharyngium* we started to observe a bizarre developmental malformation, that we nicknamed double head worms. Importantly, those animals, occurred spontaneously in our cultures, and were never very numerous – the Petri dish with hundreds of normal worms would yield less than 20 individuals, often just a few at one time. The double head worms look similar to the individuals in advanced stage of paratomy, but in addition to the anterior-facing head of the second zooid, they also have another, posterior-facing head, present at the posterior pole of the first zooid, just in front of the fission zone (Fig. 1 j–l). When we inspected details of the middle heads, we could observe that all the structures in their pharynges and brains have reversed polarity when compared to the anterior and posterior heads in the chain (Fig. 1m–v), however, they did not show any obvious malformations of the head structures. For instance, we could detect the well-developed, yet mirrored, pharyngeal musculature, including circular, longitudinal, dorsal and radial muscles (Fig. 1o, p). Similarly, the main serotoninergic structures (dorsal, and lateral perikarya, brain commissure) were present, albeit rotated by 180° (Fig. 1u). This apparently normal organization of the head tissues in the middle head of the double head worms suggests that these ectopic heads might be fully functional.

Next, we set out to investigate whether the surplus heads develop instead of the tail during paratomy, or whether they originate through the duplication of the entire A-P axis of the frontal zooids. To distinguish between those two scenarios, we needed to know whether the anterior zooid has any tail tissue located in the middle of its body (that would hint at the duplication of the A-P axis), or whether the tail identity is absent, and replaced by the head tissue instead. In contrast to the head, that can readily be distinguished based on the morphology, the tail of *Stenostomum* is an elusive body region, that does not harbor many distinct structures. However, a recently published singe-cell atlas of *S. brevipharyngium*, reported specialized hindgut cells, located in the most posterior section of the gut, that could be used as markers of the posterior identity, at least within digestive system [22]. Therefore, to test for the relation of head and tail structures in asexual and double head worms we studied the expression of brain marker, *MEF2C*, and hindgut marker, *SLC26A2*, in the animals at the advanced paratomy and with the double head phenotype.

In the asexual worms, both heads express *MEF2C* and both tails express *SLC26A2* (Fig. 2a), and expression of both markers is strongly limited to their respective territories. In the fission zone, the *SLC26A2*^*+*^ cells of the newly forming tail of anterior zooid are adjacent to the *MEF2C*^*+*^ brain cells of the second zooid (Fig. 2b–d). In the double head worms, both anterior and posterior brains of the first zooid, as well as the brain of the second zooid are expressing *MEF2C* (Fig. 2e), while *SLC26A2* is expressed in the posterior part of the second zooid (Fig. 2f). In the fission zone, two clusters of the *MEF2C*^*+*^ brain cells are located next to each other and the *SLC26A2*^*+*^ cells are missing (Fig. 2g–i). The tail marker is generally absent from the anterior zooid (Fig. 2e), although some ectopic *SLC26A2*^*+*^cells can be present next to the pharynx of the posterior-facing middle head (arrowhead, Fig. 2h–i). Generally, these expression patterns indicate that the double head worms emerge through the development of new posterior-facing head, instead of the tail, during the paratomy. Consequently, the anterior zooid of the double head worms is altogether devoid of the tail identity.

**Fig. 2.**
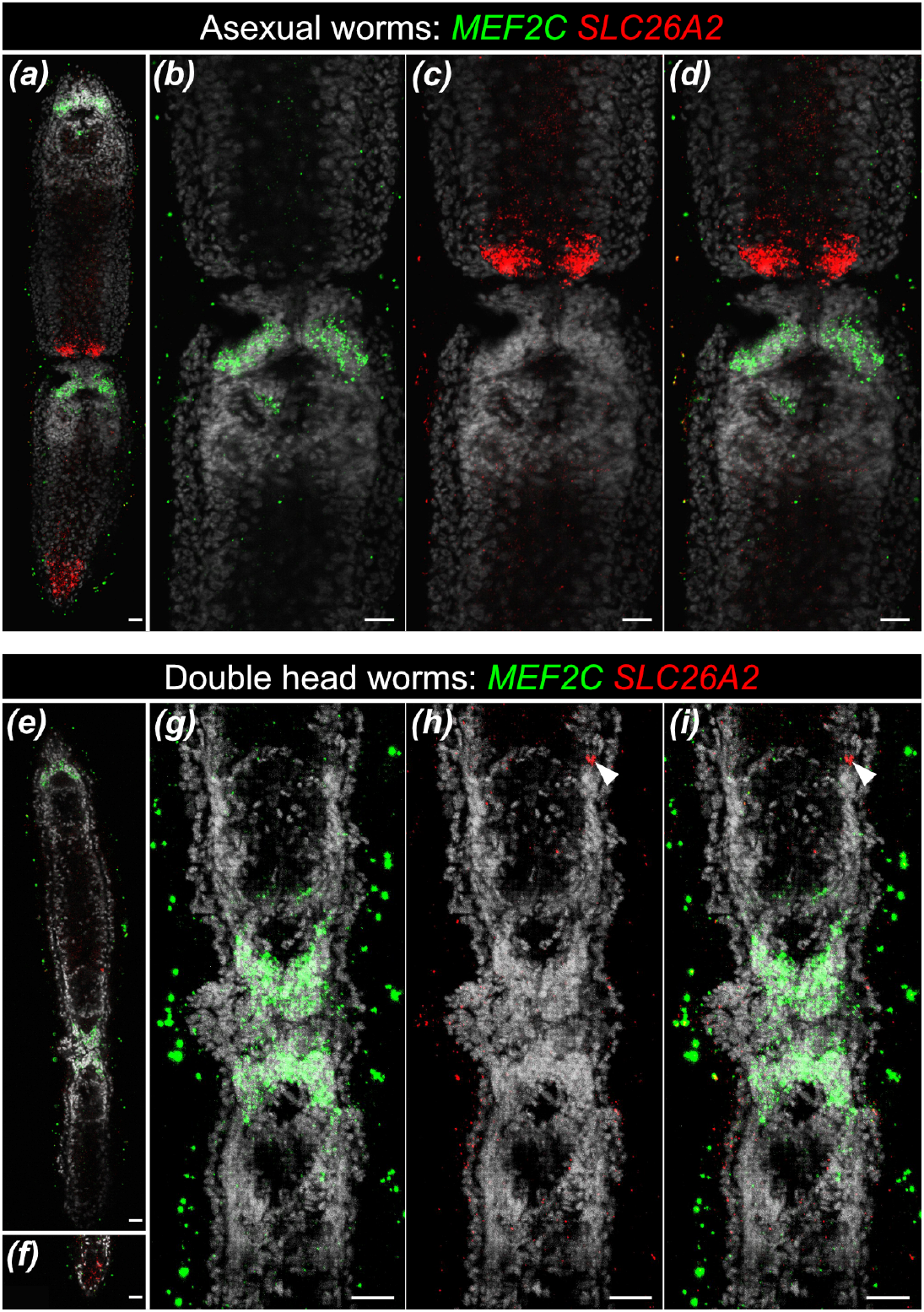
Comparison of the brain and hindgut markers expression between wild type asexual and double head *Stenostomum brevipharyngium*. **(a)** Gene expression in the worm at the advanced stage of paratomy. **(c–d)** Detailed gene expression in the fission zone of the asexual worm. **(e)** Gene expression of in the double head worm. **(f)** Gene expression in the posterior part of the posterior zooid. **(g–i)** Detailed gene expression in the fission zone of the double head worm. White arrowheads indicate ectopic *SLC26A2*^+^ cell located in the pharyngeal region. Scale bars indicate 10 μm.

### Double heads are not heritable on the organismal level

Is the double head phenotype related with some mutation, or is it a pathological condition that depends on some non-heritable factors? To address this issue, we performed set of experiments, in which we assess the heritability of the double head phenotype.

In the first experimental setup we cut six double head worms in three sections (Fig3a). An anterior worm encompassed the most anterior head and half of the first trunk. The middle worm contained the posterior-facing head and the posterior half of the first trunk. The posterior worm originated by separation of the second zooid of the chain. Additionally, we also took six normal worms, without double head phenotype, as a control group. Since anterior and middle worms were missing posterior identities (see above), we also cut off the most posterior tail tips of the posterior and control worms. First, we scored the status of regenerates at 4 days post amputation (dpa), checking whether the animals survived and what is their regenerative and reproductive status (Fig. 3b). Among control animals all worms show regular morphology and five were already engaged in the paratomy. Although single worms died or were dying among both anterior and posterior individuals, the remaining were healthy, did not develop ectopic heads, and half of them already shown symptoms of normal asexual reproduction, generally exhibiting the same outcome as control animals. The only group that significantly differed from the control worms were middle worms, among which one individual was dead, two were in advanced degeneration (resorption of the head, shrinking of the body) and three exhibited retarded regeneration. The latter animals were smaller than the control worms and either had heads looking like early regenerative stage (two individuals) or additionally had slight bumps on the body (a single individual), but they were all single headed. Altogether, we did not observe formation of ectopic heads in any of the worms originating from double head ancestors during regeneration.

**Fig. 3.**
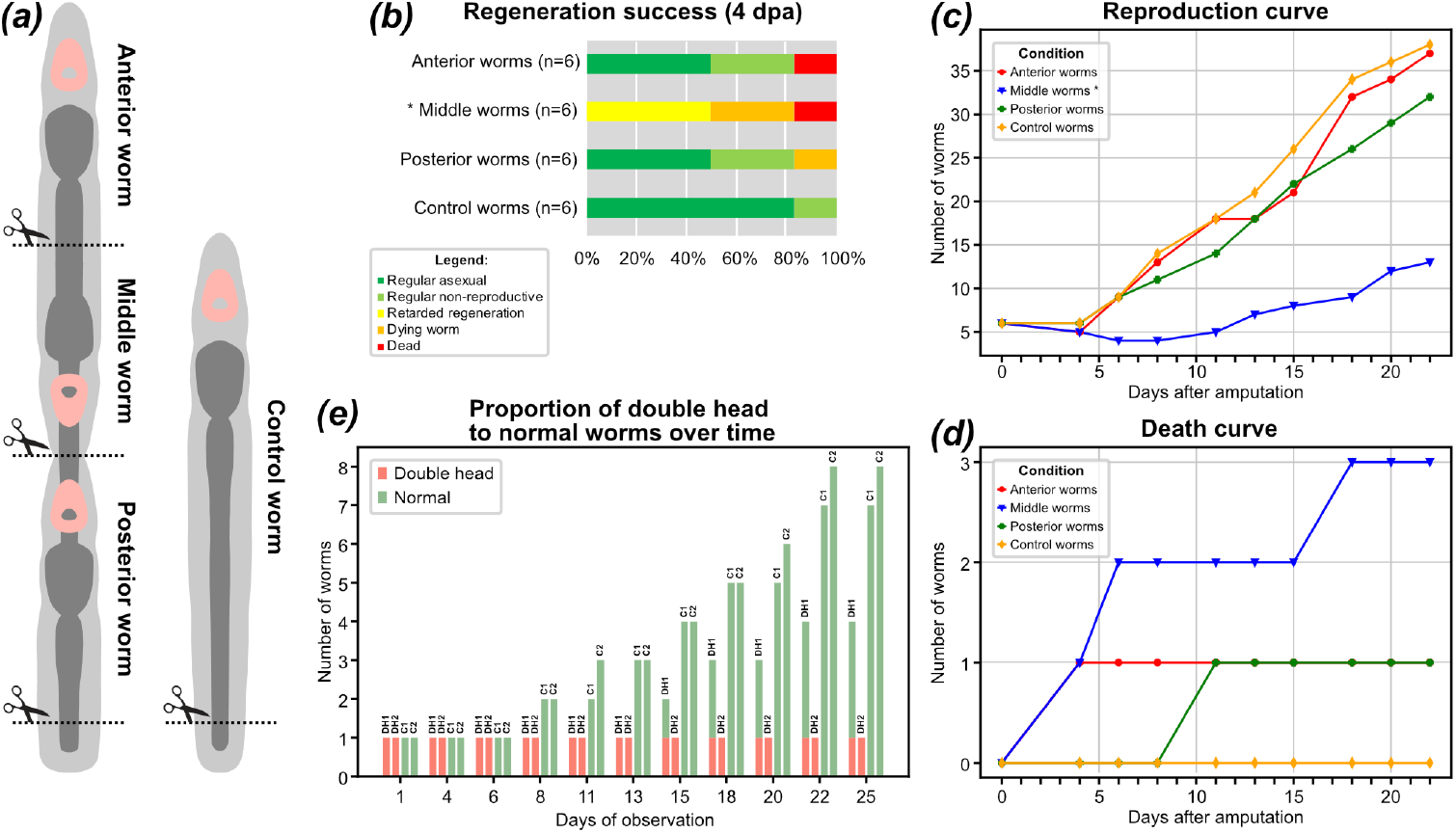
Experiments on the regeneration competence of different sections of the double head worms and heredity of the double head phenotype. **(a)** Schematic drawing of the experimental amputations. **(b)** Regeneration success at 4 days post amputation among experimental groups. **(c)** Reproduction curves of experimental groups. **(d)** Death curves of experimental groups. **(e)** Change in the proportion of double head to normal worms among the descendants of two double head (DH1 and DH2) and two control (C1 and C2) worms. Asterisks indicate significant differences from controls (p-value=0.00512) as inferred with the two-sided Mann–Whitney U test.

Next, we observed the animals that survived the regeneration over the course of following 18 days, up to 22 dpa (Table S1). The anterior and posterior worms were dividing at the same rate as control animals (Fig. 3c) and show low overall mortality rate (one animal in each group, Fig. 3d). Three worms died among middle worms (Fig. 3d) and the remaining three resumed asexual reproduction between days 8 and 15, when animals from the other groups underwent already 2-3 divisions (Table S1). In consequence, middle worms showed significantly delayed reproduction rate when compared to the other experimental groups or controls (Fig. 3c). Altogether we observed 32 divisions among both anterior and control worms, 10 divisions among middle worms, and 27 divisions among posterior worms (Table S1). Again, we did not observe any double head animals during those fissions in any of the progeny of the initial six double head individuals or controls.

In the above-described treatments, the double head worms have been cut, which might influence transmission of the phenotype to the next generation. Therefore, we set another experiment to observe the heritability of the double head phenotype in the intact animals. We took two double head and two control worms and culture them for 25 days, scoring the number of double head and normal individuals over time (Fig. 3e). While one of the double heads remained arrested in its asexual reproduction, the other divided after two weeks of observation, giving rise to a double head maternal worm and a new, normal individual. This normal animal kept dividing and did not produce the double heads, echoing the results of the first experiment.

Altogether those experiments showed that double head phenotype that we observed in S. *brevipharyngium* is not heritable on the organismal level, so it cannot be attributed to, e.g., interindividual differences in the genetic control of the asexual developmental program. What could be other potential explanation for the formation of ectopic heads? A similar phenotype, with the animals producing head instead of tail during paratomy, has been reported by Sonneborn in 1930s in *Stenostomum incaudatum* [49, 50]. In this species, the double head worms result from the ageing of the head of the most anterior zooid [49] or exposure to lead acetate [50], and represented only a small fraction of a broader spectrum of developmental malformations observed with high prevalence under those conditions. Ageing is unlikely a cause of double heads that we observed, taking into account that in our first experiment the most anterior (the oldest) head pieces of the double head worms resume asexual reproduction and their progeny did not show any malformations. Also, the laboratory cultures of *S. brevipharyngium* have been kept in captivity for more than 25 years [43] (six years by the last author), and the double head phenotype has not been observed thus far, in contrast to what would be expected from the physiologically occurring symptoms of ageing. Double head worms were present in a regular culture at low prevalence and most worms in the same culture performed unaltered paratomy, therefore an unknown toxic environmental factor is also an improbable reason for this developmental oddity. As asexual reproduction in *Stenostomum* relies on the presence of ever-dividing adult pluripotent stem cells [14, 22, 27], the observed patterns of the occurrence of double head phenotypes might be explained by the mutation at the level of single stem cells. In flatworms the morphogen gradients along A-P axis pattern stem cells differentiation to the identities specific to particular body regions [29–34]. The doble head *Stenostomum* might carry chimeric populations of stem cells that include a stem cell line that shows faulty response to those gradients. In such a case, the double head phenotype would only manifest when the stem cells originating from the mutated line are present in the fission zone and contribute to the formation of the posterior end of the anterior zooid during paratomy.

### Double heads allow reversal of the body axis polarity

In our first experiment we observed three animals with the originally posterior-facing ectopic heads that survived bisection and established viable populations through asexual reproduction. Do those worms use the ectopic head as their new anterior pole or did they resorb it, regenerate the new one at the anterior pole and thus regulate their aberrant development? To investigate this matter, we conducted another experiment, this time focusing only on the middle, posterior-facing worm and a shorter timescale (Fig. 4a). We cut five double head worms in the same way as in the first experiment, and kept the middle worms for one week in clean medium to let them regenerate. Over this time, we observed every individual at 19, 43, 67 and 93 hours post amputation (hpa), a time that is needed in *S. brevipharyngium* for complete regeneration [22, 43]. On the 7dpa the worms were fed and then observed for a week and scored for asexual reproduction. During the initial regeneration, three of the worms retained the posterior-facing heads (Fig. 4b, c) and did not grow new head structures until they engaged in paratomy, during which the new head structures were forming normally at the fission zone, facing into same direction as the first head. Two worms retained the posterior-facing heads for ca. 20hpa but later began resorbing the ectopic head (Fig. 4b, d). One of those individuals died within 2 weeks post amputation, while the other regenerated a head (although we could not determine at which end) and resumed normal asexual reproduction (Fig. 4b).

**Fig. 4.**
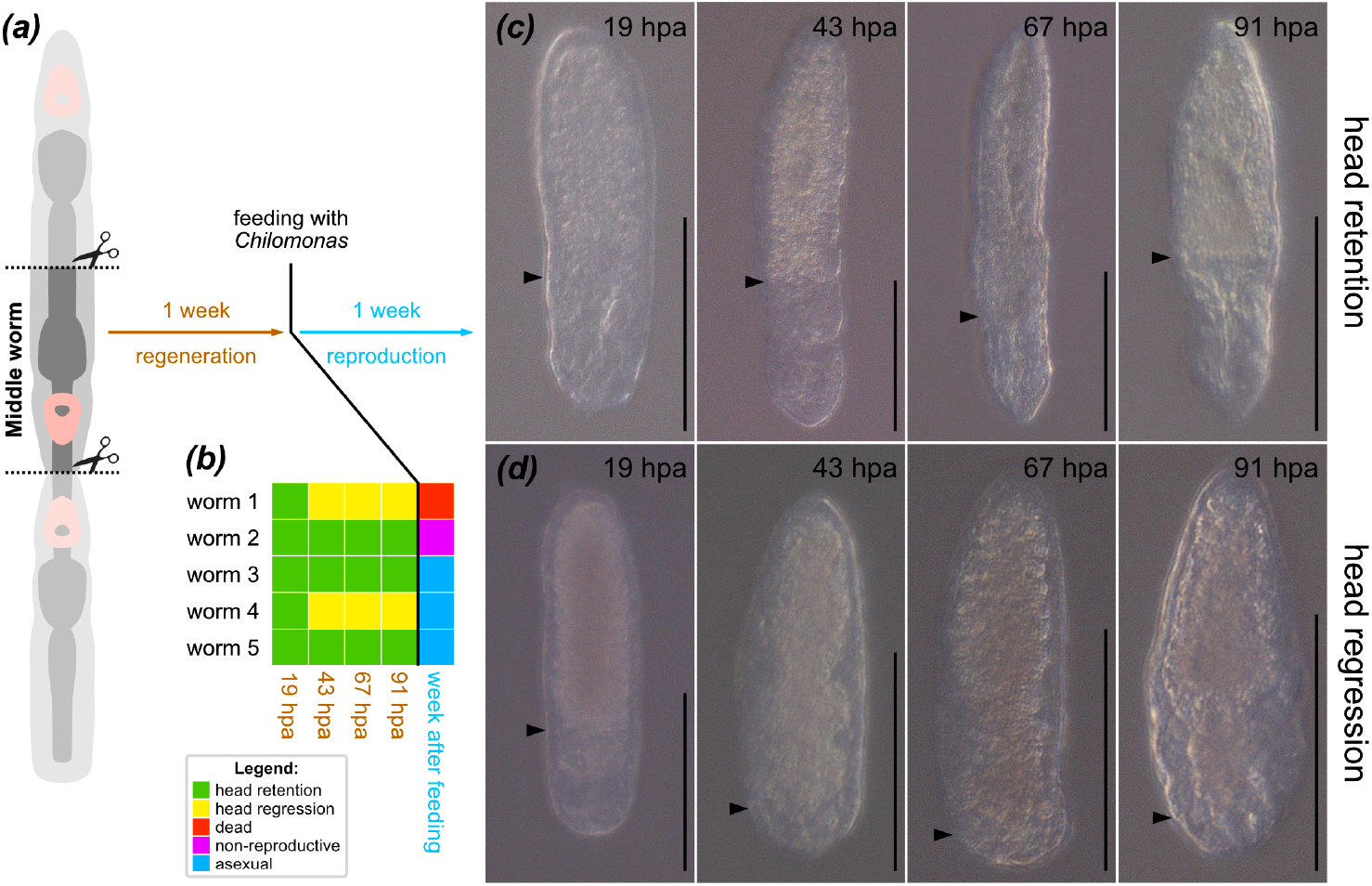
Reversal of the body axis polarity in the regenerating middle zooid of the double head worms. **(a)** Experimental design. **(b)** Regenerative and reproductive status of the worms during the experiment. **(c)** Morphological time course of head retention in the middle zooid. **(d)** Morphological time course of head regression in the middle zooid. Black arrowheads indicate the most posterior extent of the gut tissue. Scale bars indicate 100 μm.

Notably, the animals that retained the ectopic head underwent a complete 180° reversal of their A-P body axis. While their brain and pharyngeal tissues originated during the paratomy with the reversed orientation, those animals also retained trunk tissues originating from the ancestral un-inverted animal, including some that exhibit differentiation along A-P axis – e.g., body-wall musculature, protonephridium, or gut [22, 51, 52]. The fact that the worms were able to resume normal physiology, despite reversal of those vital organ systems in relation to their heads, points towards extreme physiological plasticity of their body plan. Such flexibility might be related either to relative simplicity of their organs or to the ability to dynamically remodel the tissues, due to the presence of the pluripotent stem cells [22, 27].

## Conclusions

We report here a rare double head phenotype in *Stenostomum* flatworms, that originates through the erroneous, yet spontaneous, formation of head tissues, instead of tail, during the asexual process of paratomy. While formation of the double heads in *Stenostomum* has previously been attributed to aging [49] and environmental toxicity [50], our findings suggest that it may also result from another, yet undetermined, causes – possibly linked to chimerism in adult pluripotent stem cells or stochastic perturbances of the A-P patterning molecular systems. The unique context of paratomy, and excellent regenerative capabilities of *Stenostomum* allow reversal of the A-P body polarity in the individuals originating from the pieces with the ectopic, posterior-facing heads. A similar phenomenon (ectopic head formation, followed by regeneration of a normal posterior structures on the opposite end) has been reported in some animals undergoing architomy, e.g., cnidarians [6] or planarians [37–40]. In case of a planarian *Planaria maculata* double head formation, and the following reversal of the body axis, require two rounds of cutting and regeneration (including generation of very narrow pieces, that hardly contain any tissues) [40]. In consequence, the individuals with the reversed polarity do not contain much of the tissues from the original, un-inverted worm. In case of *Stenostomum*, the double heads occur spontaneously, and contain large fragments of the regular animals, that are inherited by the worms with reversed polarity, similarly to what has been reported for a planarian *Girardia dorotocephala* [37]. We hypothesize, that developmental plasticity, driven by the presence of adult pluripotent stem cells and continuous tissue renewal [21–23], allows flatworms to survive such a dramatic developmental event.

*Stenostomum* has lately regained interest among biologists studying regeneration, asexual reproduction and stem cell biology, as an emerging and intriguing model, that in many respects differs from the other flatworms [14, 22, 43, 53]. In recent years, an increasing number of molecular tools have been developed to study these peculiar worms [14, 22, 43, 54], but our understanding of their basic biology still remains limited. The phenomenon that we report here is a prime example of how little we know about the developmental capacities of *Stenostomum*.

## Declarations

### Ethics

This work did not require ethical approval from a human subject or animal welfare committee.

### Data accessibility

The data analyzed during this study are included in this article and its supplementary materials.

## Authors’ contributions

KT discovered and collected double head worms, maintained the animal cultures and participated in documentation of the regeneration process. LG designed and conducted experiments and stainings, analyzed the data, prepared figures and drafted the manuscript. Both authors read and accepted the final version of the manuscript.

## Acknowledgments

We would like to thank Bohdan Paterczyk from the Imaging Laboratory, Faculty of Biology, University of Warsaw and Weronika Łaska, from the Institute of Evolutionary Biology, Faculty of Biology, University of Warsaw for their help with confocal and light microscopy.

